# Captivity-induced behaviour and spatial learning abilities in an enigmatic, aquifer-dwelling blind eel, *Rakthamichthys digressus*

**DOI:** 10.1101/2021.06.12.448170

**Authors:** Tanvi Vasan, Prantik Das, Vishwanath Varma, Anjani Tiwari, Archana Prakash, Devika Manilal, Liju Thomas, C. P. Arjun, Siby Philip, Rajeev Raghavan, V.V. Binoy

## Abstract

We investigated the impact of captive life on behaviour and learning abilities in an enigmatic, aquifer-dwelling blind eel, *Rakthamichthys digressus*. Of eight major behavioural traits related to exploration and activity in a novel arena, four were significantly altered by life in captivity. While the startle response upon introduction into the arena and overall swimming away from the walls increased after captivity, inactivity exhibited immediately after the startle and the reaction to an external disturbance decreased. We also observed behavioural syndromes between ‘startle responses’ and ‘horizontal wall following’, and between ‘overall activity’ and ‘vertical wall following’; however, these behavioural syndromes were not altered by maintenance in captivity. Interestingly, this blind-eel failed to learn a simple spatial task in a Y-maze apparatus. Captive-associated behavioural changes in *R. digressus* may influence their survival after reintroduction into natural habitats, and such changes must be taken into account while developing protocols for ex-situ conservation and subsequent release.

## 1. Introduction

Subterranean ecosystems, extreme environments characterized by darkness, truncated food webs and food scarcity, yet harbouring exceptional biodiversity are highly vulnerable to environmental changes [1]. Being highly restricted, and having small population sizes and low resilience, anthropogenic threats could have serious consequences on the survival of most subterranean fauna [2–3]. Though habitat loss, and contamination and overexploitation of groundwater are widely regarded as major stressors to subterranean biodiversity [4], there are also emerging threats and challenges in many regions of the world that have been poorly addressed.

A unique example, from the Western Ghats of India is ‘human-fish conflict’, where subterranean species are killed as their presence in wells is mistakenly linked to poor water quality, and some species of eels mistaken for snakes and killed on purpose [5]. Home-stead wells in this region are also cleaned annually and fish encountered during such times are also killed. To prevent this, many fish are rescued from dug-out wells and maintained in captivity until a suitable subterranean habitat is available for their release. Such captive maintenance may last for several months depending on the season of capture – e.g., rescued fish during summer may require a captive environment until their release during the subsequent monsoon.

Unlike epigean fishes in which translocation to artificial habitats are known to modify behavioural traits and syndromes [6–8], no such information is currently available on subterranean taxa. Understanding behavioural changes in captivity, and developing appropriate management protocols are critical for the success of conservation strategies such as reintroductions and translocations. This can be undertaken by altering physical properties of the environment to suit the sociobiology of the species. In many cases, it may also be required that individuals which have undergone behavioural modifications in captivity may need to be provided ‘life skill training’ [9–11] to re-organize their behavioural characters and ensure improved survival on reintroduction.

Focusing on an enigmatic, blind, synbranchid eel, *Rakthamichthys digressus*, we explore the influence of captivity on their behavioural traits and syndromes. As this species is known to inhabit a spatially complex ecosystem, i.e., narrow pores inside aquifer-bearing lateritic rocks [12], they are expected to have an excellent capacity for spatial learning. Therefore, to test this hypothesis, we assessed the spatial learning ability of *R. digressus* using a standard maze apparatus. Our study is the first to explore these behavioural aspects on blind, aquifer-dwelling eels, and provides important scientific evidence to facilitate the development of conservation strategies for subterranean fish taxa.

## 2. Materials and methods

### (a) Maintenance and husbandry

*Rakthamichthys digressus* (N=24) were collected from homestead dug-out wells as part of a rescue effort and transferred to the laboratory where they were housed in pairs in well-aerated glass aquaria (39×26×30 cm). Pieces of Poly Vinyl Chloride (PVC) pipe (diameter 2.5 cm; length 15 cm) were provided as shelters and water level was maintained at a height of 16 cm. We paired fishes slightly different in their body length (< 1 cm) to facilitate individual identification. Fish were fed with tubifex worms ad libitum in the afternoon, and uneaten food siphoned out after 30 minutes. Water temperature was maintained at 25°C and the room was maintained under a 12h:12h LD cycle. All fish were healthy until the completion of experiments, and no mortality was observed.

### (b) Experiment 1: Exploratory behaviour and syndromes

Exploratory behaviour of *R. digressus* was studied using a rectangular open-field apparatus. An aquarium (50 × 39 × 30 cm) with 2 × 2 cm grids marked on the bottom to quantify locomotor activity was the open field used in this experiment. This apparatus was filled with water up to a height of 15 cm and a Compact Fluorescent Lamp (CFL; 20W) lighted the apparatus from above. After acclimation period of seven days, subject fish were introduced individually into the middle region of the open field. After allowing six minutes for the fish to explore the arena, a small aquarium net (10.16 × 7.62 cm, handle length 29.21 cm and weight 38 grams) was dropped from a height of 15 cm at a point 10 cm away from the head of the subject fish and retrieved [13]. Fish recuperated quickly and the behaviour was recorded for two minutes after dropping the net, and the fish subsequently transferred back to its home tank (Trial 1). Subject fish were tested again using the same apparatus and protocol after 45 days to examine the impact of captive life on these behaviours (Trial 2). All experiments were conducted during the day time and recorded using a Handycam (Sony HDR-CX405) fixed above the open field apparatus.

We compared syndromes between behavioural traits in trials 1 and 2 to improve our understanding of the effect of captivity on behavioural syndromes – the tight linkage between various components of exploratory behaviours.

### (c) Experiment 2: Spatial learning ability

Spatial learning ability was studied using a ‘Y-maze’ [14] made of plexiglass sheets fixed inside an aquarium (50 × 39 × 30 cm) divided into two chambers - A (7 × 39 × 30 cm) and B (43 × 39 × 30 cm). The starting arm of the maze (31 × 2 × 30 cm) was connected to the start chamber (chamber A) by a guillotine door, and both choice arms (31 × 2 × 30 cm) were placed in chamber B. One of the arms of the Y-maze was closed at the end (blocked arm), while the other led to an open area (12 × 35 × 30 cm) which the fish could explore upon entry. A 20 W CFL lit the apparatus from above. In this experiment, we used the same individual fish that were part of trials 1 and 2, after providing an interval of 7 days. Fish was introduced individually in the start chamber, and an acclimatization time of five minutes was provided before opening the guillotine door. Fish behaviour was recorded for 15 minutes, after which it was returned to its home tank. All 24 individuals were tested once daily for six consecutive days following the same protocol.

### (d) Analysis

Videos of experiments 1 and 2 were analysed using Behavioral Observation Research Interactive Software (BORIS) [15]. Eight behavioural parameters were quantified from the videos of the experiment 1 and three parameters from experiment 2 (Table S1). Analyses were carried out using Linear Mixed Modelling (LMM) using R version 3.6.1 [16]. Behavioural data (after log transformation) followed normal distribution, and were used as dependent variables in the analysis. Trial number (trial 1 refers to the assay performed in the first week after transfer to laboratory conditions, and trial 2 to that performed after 45 days in captivity) was considered an independent variable. Significant effects of trial number suggest that dependent variable (behavioural traits) was altered as a consequence of captive life. Identity of individual fish was used as the random factor in these analyses. For examining changes in behavioural syndromes after captivity, we examined effect of the interaction term between behavioural traits and trial number in the linear mixed model. We performed this analysis by comparing models with, and without such interaction terms. Models with lowest AIC values were considered the best fits, and models with *Δ*lAIC > 2 significantly poorer fits. If models with interaction between behavioural traits and trial number were significantly better fits than those without, then we inferred that behavioural syndrome was altered by captivity. In the case of experiments examining the spatial learning ability, data on selection of the arm of Y maze, a binary choice situation, was modelled as a binomial distribution. Hence, Generalized Linear Mixed Models (GLMM) was used. The packages used were ‘ggplot2’, ‘lmerTest’, and ‘lme4’ [16]

## 3. Results

### (a) Effect of captivity on behavioural traits and syndromes

Life in captivity affected certain behaviour traits such as sudden movements upon introduction into the novel arena (startle response, β = 10.05, p = 0.05) and ‘swimming away from the walls’ (β = 8.60, p = 0.02) which significantly increased, and ‘duration of rapid swimming’ in response to dropping the net (reaction, β = -7.89, p = 0.001) and ‘total time spent in rest during exploration’ (rest, β = -7.73, p = 0.002, Figure 1a; Table 1) which decreased. These results are substantiated by the fact that models without trial as an independent factor were poorer fits for these traits (*Δ*lAIC > 2; Table 2). However, duration of inactivity bout immediately after startle (‘inactivity after startle’ β = -4.54, p = 0.06), ‘moving along the wall of the apparatus towards the water surface’ (vertical wall following, β = 0.91, p = 0.83, Figure 1a), and ‘swimming along the wall parallel to it’ (horizontal wall following behaviour, β = -1.55, p = 0.86, Figure 1a) and ‘total activity’ (β = 7.17, p = 0.40, Figure 1a) were not affected by captivity. We did not observe any significant difference in the fit of models of these behavioural traits with, and without trial, as independent factor (*Δ*lAIC < 2; Table 2).

**Figure 1.**
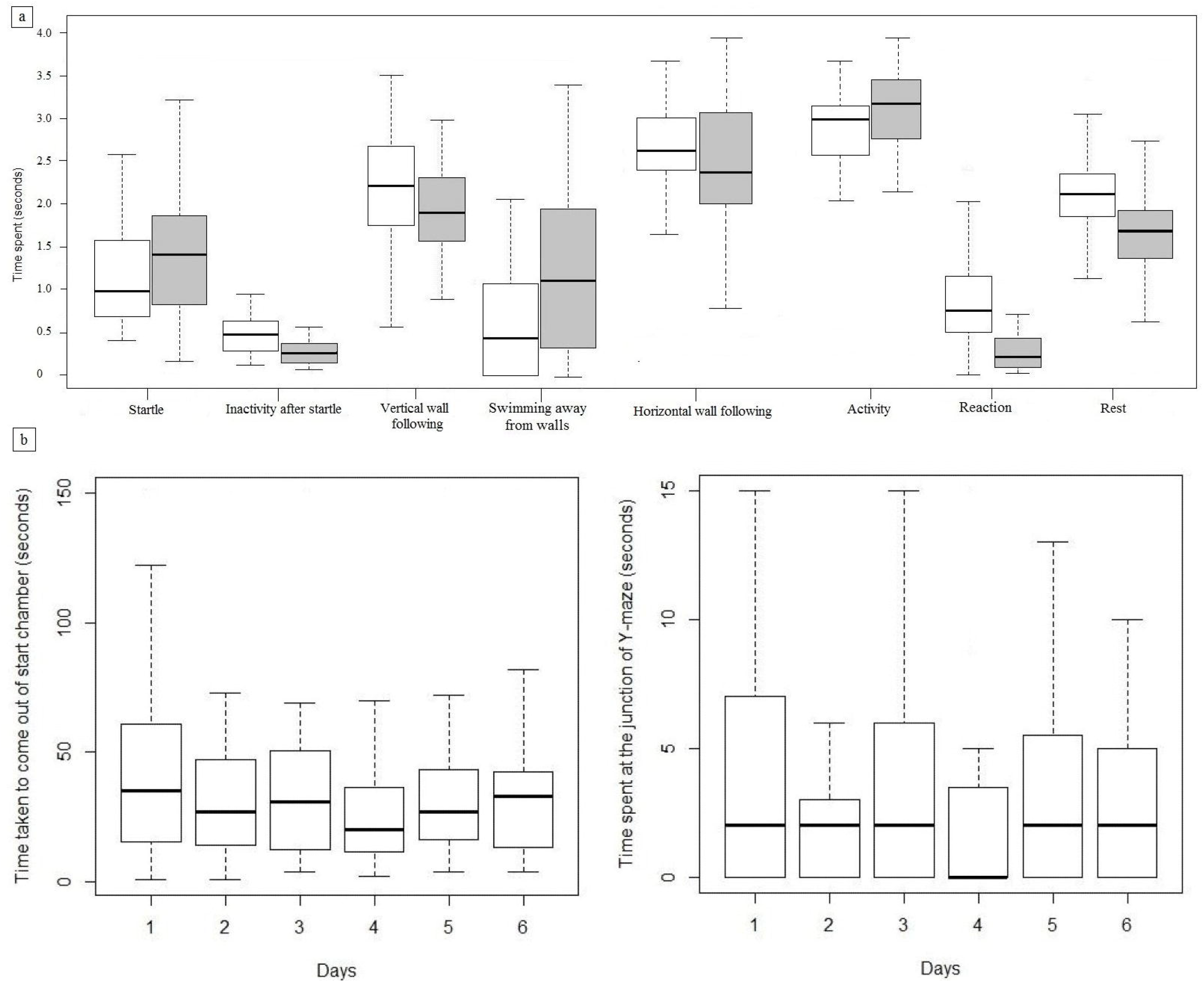
(a) Influence of captivity on various elements of exploratory behaviour of *Rakthamichthys digressus*; open and shaded boxplots represent Trial 1 (5^th^ day) and Trial 2 (45^th^ day) respectively. (b) Latency to enter the maze and time spent in the junction of the choice arms by the test fish during spatial learning experiment conducted using Y maze.

**Table 1.** Influence of 45-day captivity on various elements of exploratory behaviour exhibited by *Rakthamichthys digressus* (Experiment 1). Experiment 2 represents the effect of repeated exposure to Y maze for five consecutive days on the spatial learning ability in this species. Statistics used were LMM in all cases except arm choice (GLMM), which was a binary data. *= P<0.05,**= P<0.01, ***= P<0.0001.

| S. No. | Behavioural parameters | $\beta$ | p |
| --- | --- | --- | --- |
| **Experiment 1 (Exploratory behaviour)** |  |  |  |
| 1. | Startle | 10.05 | 0.05* |
| 2. | Inactivity after startle | -4.54 | 0.06 |
| 3. | Rest | -7.73 | 0.002** |
| 4. | Swimming away from walls | 8.60 | 0.02* |
| 5. | Horizontal wall following | -1.55 | 0.86 |
| 6. | Vertical wall following | 0.91 | 0.83 |
| 7. | Activity | 7.17 | 0.40 |
| 8. | Reaction | -7.89 | 0.001** |
| **Experiment 2 (Spatial learning)** |  |  |  |
| 9. | Maze entry | -1.73 | 0.38 |
| 10. | Time spent at the junction | -0.19 | 0.70 |
| 11. | Arm choice | -0.02 | 0.83 |

**Table 2.** Model comparisons of linear mixed models (LMM) with various behavioural traits of *Rakthamichthys digressus* studied during the ‘Trials 1 and 2’ of experiment 1. Each row shows the results of different models with degrees of freedom and difference in AIC (Akaike Information Criterion) values from the best-fitted model. The models are ordered with increasing AIC values.

| Response Variable | Model | df | ΔAIC |
| --- | --- | --- | --- |
| Startle | HorizontalWallFollowing * Trial + (1\|FishID) | 6 | 1.93 |
|  | HorizontalWallFollowing + Trial + (1\|FishID) | 5 | 0 |
|  | HorizontalWallFollowing + (1\|FishID) | 4 | 2.58 |
|  | Trial + (1\|FishID) | 4 | 10.11 |
|  | 1 + (1\|FishID) | 3 | 21.26 |
| Activity | VerticalWallFollowing * Trial + (1\|FishID) | 6 | 1.53 |
|  | VerticalWallFollowing + Trial + (1\|FishID) | 5 | 0.98 |
|  | VerticalWallFollowing + (1\|FishID) | 4 | 0 |
|  | Trial + (1\|FishID) | 4 | 24.99 |
|  | 1 + (1\|FishID) | 3 | 18.88 |

In terms of behavioural syndromes, the ‘startle response’ was negatively correlated with ‘horizontal wall following behaviour’ (β = -0.24, p < 0.0001; Figure 2a). Full Linear Mixed Models of ‘startle response’ as response variable with interactions between ‘wall following behaviour’ and trial revealed that the model without interaction was the best fit for ‘startle response’ (*Δ*lAIC = 1.93 for the model with interaction; Table 2). This suggests that the behavioural syndrome between ‘startle response’ and ‘swimming along walls’ was not influenced by 45 days of captivity. Similarly positive correlation noted between ‘vertical wall following’ and ‘activity’ (β = 1.33, p < 0.0001; Figure 2b) was also found to be uninfluenced by the captive life. No linkage was observed between any other behaviour parameters studied.

**Figure 2.**
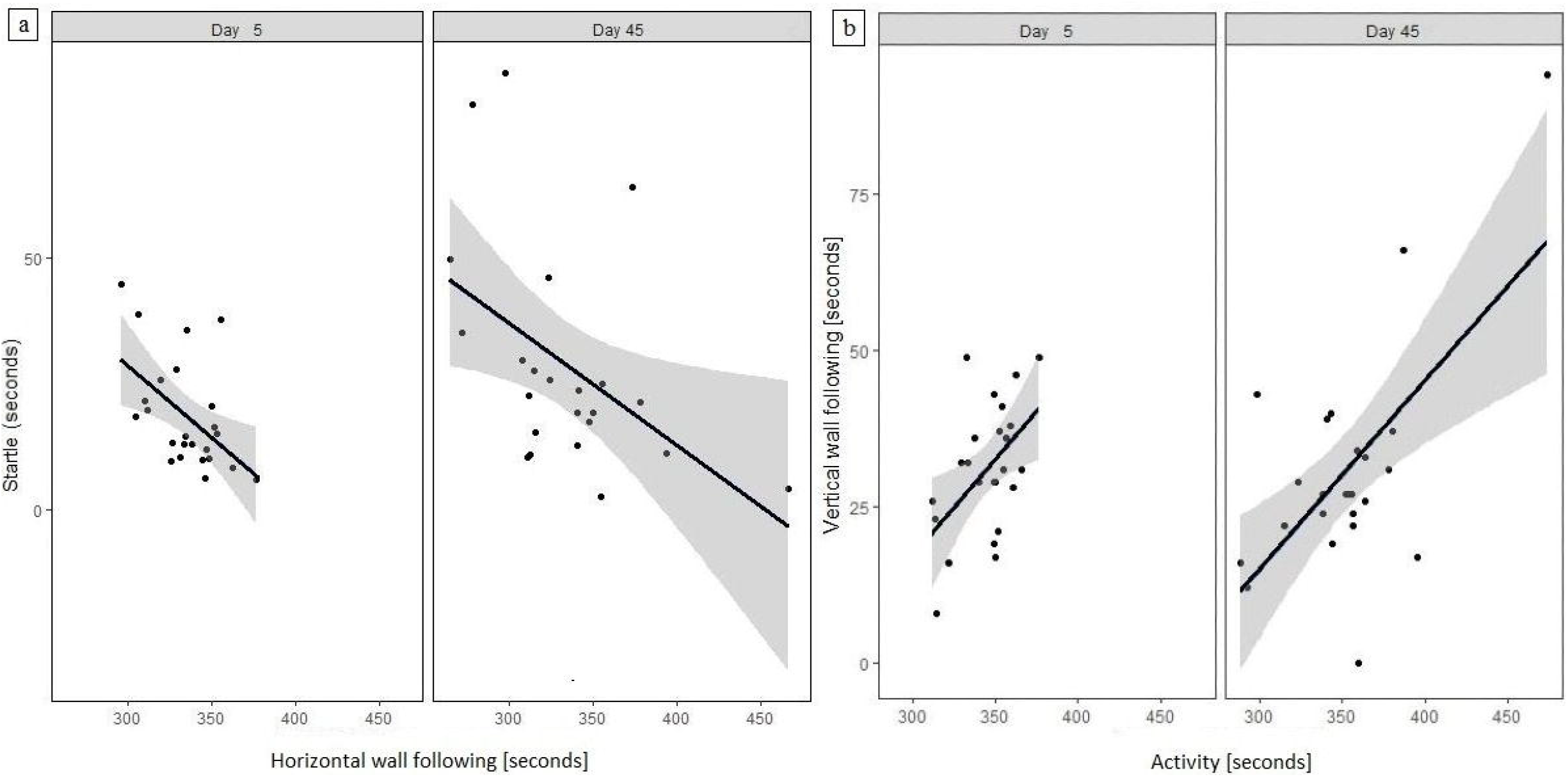
Influence of captivity on the correlations between behaviour traits (a) startle and horizontal wall following and (b) vertical wall following and activity in *R. digressus*. Each dot represents individual fish (n = 24) and the grey shading represents 95% confidence intervals.

### (b) Spatial learning

No significant change was observed in the ‘latency to leave the start chamber’ (LMM: β = - 1.73, p = 0.34; Table 1; Figure 1b), ‘time spent at junction of the Y-maze’ before choosing an arm (LMM: β = -0.19, p = 0.68; Figure 1b), and ‘selecting one arm over the other’ (GLMM-binomial: β = -0.02, p = 0.83), over repeated testing conducted for six days. This revealed that *R. digressus* did not develop preference towards any choice arm, indicating that it failed to learn this simple spatial task.

## 4. Discussion

Temporary translocation of threatened fish from their natural habitat to captivity, and reintroduction during favourable times is a strategy for avoiding permanent loss of individuals or fragmentation of populations during harsh seasons [17,18]. Translocation and subsequent life in captivity for 45 days significantly increased the ‘startle response’, ‘time spent swimming away from the walls’, and ‘rest taken during the exploration’ in *R. digressus*. Meanwhile, the reaction towards a fear inducing stimulus became weaker. Startle response exhibited by a fish in open field is attributed to fear, or handling by humans [19,20]. However, separating the effect of these two parameters on the enhanced startle exhibited by captive *R. digressus* is difficult [20]. Another behavioural expression of fear, i.e., ‘reaction,’ which diminished in captivity suggests that handling by humans may be potentially promoting startle response.

Blind subterranean fishes which possess minimal sensory range are highly dependent on tactile stimuli and lateral line sense organs to perceive their environment [21,22]. This may explain why they follow the walls of the novel environment so that their snouts bestowed with lateral line sensory organs could receive more sensory stimulus. Previous research has demonstrated that impairment of the lateral line system is associated with reduced ‘wall following behaviour’ [23]. Though the exact reason for the reduction in ‘time spent near wall’ could not be understood, diameter of snout of the eel (where lateral line pore system is pronounced [24]) was observed to have reduced in captivity (results not shown).

Moving individuals from their natural habitat to captivity has influenced behavioural syndromes in many epigean fishes [7,8]. In *R. digressus*, a negative correlation between ‘startle response’ and ‘horizontal wall following’, and a positive linkage between ‘activity’ and the ‘vertical wall following’ was observed. Individuals that exhibit higher levels of startle response due to higher neophobia spend more time near the sides of the open field in species ranging from rodents to fishes [25–27]. Active *R. digressus*, spending more time in ‘vertical wall following’ could either be due to the inability to recognize the presence of wall [23], or searching for biologically significant resources similar to its function in natural habitats. Although boldness - propensity to take a risky decision is positively correlated with activity in many epigean fishes [28,29], ‘startle response’, duration of the ‘inactivity after the startle’, and ‘reaction’, which are potential indirect measures of boldness [30,31] failed to show any association with activity in *R. digressus*. Furthermore, life in captivity for 45 days neither changed any existing behavioural syndromes, nor generated new associations between behaviour traits in this species.

In spite of a short sensory range, blind fishes have been known to learn spatial properties of their environment [32,33]. However, in *R. digressus*, latency to leave the start chamber did not change and no preference was developed towards any of the choice arms of Y maze indicating the inability to learn this simple spatial task. This lack of learning may be due to the negative effects of captive conditions, short duration of exposure (15 minutes for 5 consecutive days) to the maze or neither of the choice arms being a strong reward [34]. Hence, analysing the spatial learning ability in *R. digressus* immediately after collecting from the natural habitat and providing more time to familiarise with the spatial properties of the apparatus is essential to distinguish between whether this inability is a species-specific characteristic or the consequence of captive life.

Captivity altered certain behavioural traits such as ‘increased startle response’ and ‘swimming away from the wall’, and ‘reduced fear response’ in *R. digressus*, which may increase its predation risk when reintroduced into its natural habitat. Hence, to mitigate such adverse effects of captive life, increasing complexity of artificial habitat [9], soft release [35], and life skill training protocols [11] should be considered for improving the success of their reintroduction.

## Acknowledgements

The senior author is grateful to the Rufford Foundation UK for Small Grant (No 17135-1) for funding, and local communities in Kannur District of Kerala for their help and support in the field.

## Notes

### Competing Interest Statement

The authors have declared no competing interest.

