## Supplementary material for "Captivity-induced behaviour and spatial learning abilities in an enigmatic, aquifer-dwelling blind eel, *Rakthamichthys digressus*": Table S1

**Supplementary Table**

**Table S1.** Behavioural parameters quantified in Experiment 1 (exploratory behaviour) and Experiment 2 (spatial learning).

**S. No. Behavioural parameters	Description**

**Experiment 1 (Exploratory behaviour)**

1. Startle	Quick and abrupt movement exhibited by the

subject fish immediately after introduction into

the open field

2. Inactivity after startle	Latency to restart exploration after the short

inactive state post startle response

3. Rest	Time spent in rest while exploring the open field
4. Swimming away from walls	Swimming at least 1cm away from the walls

of the open field

5. Horizontal wall following  Swimming parallel to the side walls of the

tank

6. Vertical wall following   Moving vertically towards the water surface

along the sidewalls of the open field apparatus

7. Activity	Total time spent actively moving in the open

field

8. Reaction	Abrupt movements exhibited in response to the

fishing net falling into the open field

**Experiment 2 (Spatial learning)**

9. Maze entry	Latency to move from start chamber to  the start

arm of the Y-maze after the guillotine door is

open

10. Time spent at the junction	Time spent at the junction between start and

choice arms of the maze before entering into one

of the choice arm

11. Arm choice	Entering into one of the choice arm
